# *Cis*-regulatory divergence in gene expression between two thermally divergent yeast species

**DOI:** 10.1101/086785

**Authors:** Xueying C. Li, Justin C. Fay

## Abstract

Gene regulation is a ubiquitous mechanism by which organisms respond to their environment. While organisms are often found to be adapted to the environments they experience, the role of gene regulation in environmental adaptation is not often known. In this study, we examine divergence in *cis*-regulatory effects between two *Saccharomyces* species, *S. cerevisiae* and *S. uvarum*, that have substantially diverged in their thermal growth profile. We measured allele specific expression (ASE) in the species’ hybrid at three temperatures, the highest of which is lethal to *S. uvarum* but not the hybrid or *S. cerevisiae*. We find that *S. uvarum* alleles can be expressed at the same level as *S. cerevisiae* alleles at high temperature and most *cis*-acting differences in gene expression are not dependent on temperature. While a small set of 136 genes show temperature-dependent ASE, we find no indication that signatures of directional *cis*-regulatory evolution are associated with temperature. Within promoter regions we find binding sites enriched upstream of temperature responsive genes, but only weak correlations between binding site and expression divergence. Our results indicate that temperature divergence between *S. cerevisiae* and *S. uvarum* has not caused widespread divergence in *cis*-regulatory activity, but point to a small subset of genes where the species’ alleles show differences in magnitude or opposite responses to temperature. The difficulty of explaining divergence in *cis*-regulatory sequences with models of transcription factor binding sites and nucleosome positioning highlights the importance of identifying mutations that underlie *cis*-regulatory divergence between species.

## Introduction

Changes in gene regulation are thought to play an important role in evolution (Carroll 2000). Regulatory change may be of particular importance to morphological evolution where tissue specific changes and co-option of existing pathways can modulate essential and conserved developmental pathways without a cost imposed by more pleiotropic changes in protein structure. Indeed, many examples illustrate this view and there is a strong tendency for *cis*-acting changes in gene expression to underlie morphological evolution between species (Stern and Orgogozo 2008).

However, gene regulation is also critical to responding to environmental changes and all organisms that have been examined exhibit diverse transcriptional responses that depend on the environmental alteration (López-maury et al. 2008). Environment-dependent gene regulation enables fine-tuning of metabolism depending on nutrient availability as well as avoiding the potential costs of constitutive expression of proteins that are beneficial in certain environments but deleterious in others. Despite the general importance of responding to changing environments, the role of gene regulation in modulating these responses between closely related species is not known and may involve structural changes in proteins whose expression is already environment-dependent.

Studies of genetic variation in gene expression within and between species have revealed an abundance of variation (reviewed in Whitehead and Crawford 2006, Zheng et al. 2011, Romero et al. 2012). When examined, a significant fraction of this variation is environment-dependent (Fay et al. 2004, Landry et al. 2006, Li et al. 2006, Smith and Kruglyak 2008, Tirosh et al. 2009, Fear et al. 2016, He et al. 2016; reviewed in Gibson 2008, Grishkevich and Yanai 2013). However, distinguishing between adaptive and neutral divergence in gene expression is challenging (Fay and Wittkopp 2008), since *trans*-acting changes can cause correlated changes in the expression of many genes and the rate of expression divergence depends on the mutation rate and effect size, which is likely gene-specific and not known for all but a few genes (Gruber et al. 2012, Yun et al. 2012, Metzger et al. 2015).

One potentially powerful means of identifying adaptive divergence in gene expression is through a sign test of directional *cis*-acting changes in gene expression measured by allele-specific expression (ASE) (Fraser 2011). By testing whether a group of functionally related or co-regulated group of genes have evolved consistently higher or lower expression levels, the test does not assume any distribution of effect sizes and more importantly is specifically targeted to identifying polygenic adaptation. Applications of this or related sign tests (Fraser et al. 2010, Naranjo et al. 2015) have revealed quite a few cases of adaptive evolution (Bullard et al. 2010, Fraser et al. 2010, Fraser et al. 2011, Fraser et al. 2012, Martin et al. 2012, Chang et al. 2013, Naranjo et al. 2015, He et al. 2016, Roop et al. 2016), some of which have been linked to organismal phenotypes. However, in only two of these studies was condition-specific divergence in gene expression examined (He et al. 2016, Roop et al. 2016), leaving open the question of how often such changes exhibit evidence for adaptive evolution. Of potential relevance, the majority (44-89%) of environment-dependent differences in gene expression have been found to be caused by *trans* rather than *cis*-acting changes in gene expression (Smith and Kruglyak 2008, Tirosh et al. 2009, Grundberg et al. 2011, Fear et al. 2016), suggesting that *trans*-acting changes in gene expression may be more important to modulating environment-dependent gene expression.

In this study, we examine allele-specific differences in expression between two *Saccharomyces* species that have diverged in their thermal growth profiles. Among the *Saccharomyces* species, the most prominent phenotypic difference is in their thermal growth profile (Gonçalves et al. 2011,Salvadó et al. 2011). The optimum growth temperature of *S. cerevisiae* and *S. paradoxus* is 29–35°C, whereas the optimum growth temperature for *S. uvarum* and *S. kudriavzevii* is 23–27°C (Salvadó et al. 2011). Furthermore, *S. cerevisiae* is able to grow at much higher temperatures (maximum 41–42°C) than *S. uvarum* (maximum 34–35°C, Gonçalves et al. 2011), while *S. uvarum* grows much better than *S. cerevisiae* at low temperature (4°C, Figure 1). Because *S. cerevisiae* × *S. uvarum* hybrids grow well at high temperature, we were able to measure *cis*-regulatory divergence in gene expression across a range of temperatures by measuring ASE in the hybrid. We use this approach to determine how ASE is influenced by temperature and specifically whether *S. uvarum* alleles are misregulated at temperatures not experienced in their native context. We find that most ASE is independent of temperature and only a small subset of genes show an allele-specific temperature response.

**Fig. 1.**
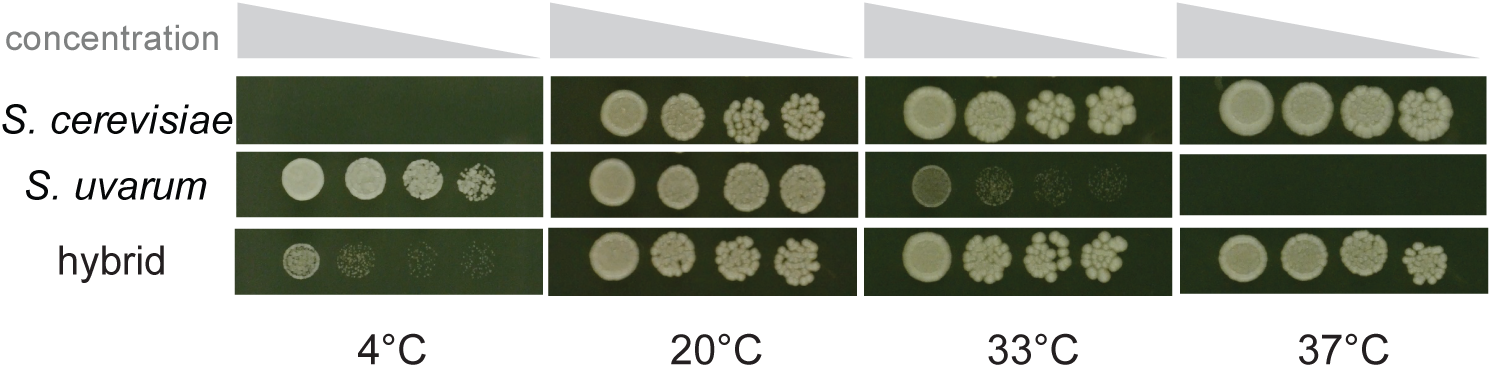
Temperature dependent growth of *S. cerevisiae, S. uvarum* and their hybrid. Growth is after 17 days at 4°C, 3 days at 20°C and 2 days at 33°C and 37°C, with platings on YPD at 1:3 serial dilutions.

## Material and methods

### Strains and RNA sequencing

A hybrid strain YJF1484 was made by crossing an *S. cerevisiae* strain YJF153 *(MATa hoΔ*∷*dsdAMX4*, derived from an oak tree isolate YPS163) and an *S. uvarum* strain YJF1450 *(MATα hoΔ*∷*NatMX*, derived from CBS7001 and provided by C. Hittinger). The hybrid was typed by PCR (Albertin et al. 2013) and found to carry *S. cerevisiae* mitochondrial DNA. A diploid *S. cerevisae* strain YJF1463 was made by crossing YJF153 *(MATa hoΔ::dsdAMX4)* and YJF154 *(MATα hoA*∷*dsdAMX4*, derived from YPS163). The diploid *S. uvarum* strain YJF2602 was made by crossing YJF1449 (*MATa hoΔ*∷*NatMX*, derived from CBS7001) and YJF1450 (*MATα hoΔ*∷*NatMX*).

Three replicate overnight cultures of the diploid hybrid YJF1484 were used to inoculate 50 ml YPD cultures (1% yeast extract, 2% peptone, 2% glucose) and incubated at either 22°C, 33°C or 37°C at 300 rpm. Cells were harvested at mid-log phase and RNA was extracted with phenol/chloroform. The nine RNA samples were enriched for mRNA by poly A purification, reverse transcribed, fragmented, ligated to indexed adaptors and sequenced on a HiSeq (1×50 bp run) at Washington University’s Genome Technology Access Center.

### Allele-specific expression differences

Reads were mapped using Bowtie2 (Langmead and Salzberg 2012) to a combined *S. cerevisiae* and *S. uvarum* genome. The YJF153 genome was generated by converting the S288c (R64-1-1) reference to YJF153 using GATK (v3.3-0) and YJF153 variants. YJF153 variants were called using GATK and 5.3 million paired-end (2×101 bp) HiSeq reads (SRX2321838). Annotations for the YJF153 genome were obtained using S288c annotations and the UCSC LiftOver tool. The YJF1450 genome and annotation files were obtained from Scannell et al. (2011). We obtained an average of 5.5 million mapped reads per sample after removing duplicate reads and reads with low mapping quality (MQ < 2). All the remaining reads were uniquely mapped as they had a higher primary than secondary alignment score (AS>XS).

Read counts for each gene were generated using HTSeq-count (Anders et al. 2015) with the default settings, which only counts reads with a mapping quality of at least 10. Species-specific counts of 5,055 orthologs were generated using previously defined one-to-one orthologs (Scannell et al. 2011). To quantify any systematic bias in read mapping we calculated the ratio of normalized *S. cerevisiae* to *S. uvarum* expression levels and found a median of 0.998, indicating no systematic read mapping bias. In our data, expression differences did not correlate with GC content (p = 0.74, linear regression), which was a concern in a previous report (Bullard et al. 2010).

Significant differences in expression were tested using a generalized linear model with a negative binomial error model (Anders et al. 2010). Using normalized read counts we tested each gene for i) temperature effects, ii) allele effects, and iii) temperature-allele interactions by dropping terms from the full model: *counts ~ allele + temperature + allele*temperature*, where *allele* and *temperature* are terms indicating the species’ allele and temperature effect and the star indicates an interaction. For score assignment in the sign test (see below), we treated data from three temperatures separately and tested each gene for allele-specific expression (ASE) at each temperature. A false discovery rate (FDR) cutoff of 0.05 was used for significance.

### Quantitative PCR

Quantitative PCR (qPCR) was used to quantify the expression of *HSP104* in the hybrid as well as both parental strains following temperature treatment. Overnight cultures were grown at 23°C, diluted to an optical density (OD600) of 0.1 in YPD for temperature treatment and grown at 10°C, 23°C and 37°C for two days, 6 hours and 5 hours respectively. The middle and high temperature cultures were shaken at 250 rpm whereas the low temperature cultures were grown without shaking. At the time of collection, the OD600 of the cultures were all within the range of 0.5 – 1.9. RNA was extracted as described above, DNase I treated (RQ1 RNase-free DNase, Promega) and cDNA was synthesized (Protoscript II Reverse Transcriptase, New England Biolabs). qPCR amplifications used the Power SYBR Green Master Mix (Thermo Fisher Scientific Inc.) and were quantified on an ABI Prism 7900HT Sequence Detection System (Applied Biosystems). Each PCR reaction was run in triplicate and one sample was removed from analysis due to a high standard error of deltaCt values (>0.4) among the three technical replicates. For each sample, expression of *HSP104* was measured relative to *ACT1* expression. Because we used allele-specific primers to distinguish *S. cerevisiae* and *S. uvarum* alleles of *HSP104*, the expression levels were corrected using the PCR efficiency of each primer sets, determined by standard curves. Genomic DNA of YJF1484 was used as a calibrator and to remove any plate-to-plate differences.

### Sign test for directional divergence

Pathways and groups of co-regulated genes were tested for directional divergence using a sign test as previously described (Bullard et al. 2010). Each gene was assigned a score 0 if the gene showed no ASE, −1 if the gene showed ASE and the *S. cerevisiae* allele was expressed higher than the *S. uvarum* allele and −1 if the gene showed ASE and the *S. cerevisiae* allele was expressed lower than the *S. uvarum* allele. Scores for all the genes in a co-regulated group (Gasch et al. 2004) were summed and tested for significant deviations from 0 by permutation resampling of scores across all 5055 genes. To correct for multiple comparisons, the false discovery rate was estimated from the permuted data across all groups. The analysis was independently applied to data from 22°C, 33°C and 37°C.

### Association with genomic features

Expression levels were associated with features of intergenic sequences, defined as sequences between annotated coding sequences. Intergenic sequences were obtained from http://www.SaccharomycesSensuStricto.org and pairwise alignments were generated using FSA (Bradley et al. 2009). Substitution rates were calculated using the HKY85 model of nucleotide substitution implemented in PAML (Yang 2007).

Transcription factor binding site scores were generated by Patser (Hertz and Stormo 1999) with 244 position weight matrix (PWM) models from YeTFasCo (expert-curated database, de Boer and Hughes 2012), using a pseudocount of 0.001. Binding site scores are the log-likelihood of observing the sequence under the motif model compared to a background model of nucleotide frequencies (G+C = 34.2% for *S. cerevisiae* and 36.3% for *S. uvarum*). For each gene we used the highest scoring binding site within its upstream intergenic region. Negative scores were set to zero. The temperature effects of *S. cerevisiae* alleles were used in the following analysis. Binding sites associated with temperature effects were identified by linear regression with the average binding site score of the two species. Mann-Whitney tests were used to assess enrichment of binding sites in temperature-responsive genes compared to genes without a temperature response. Motif models that were significant for both linear regression and Mann-Whitney tests after Holm-Bonferroni correction were considered positive hits.

Predicted nucleosome occupancy was generated by NuPoP (Xi et al. 2010), using the yeast model for both species. The average nucleosome occupancy across each promoter was used. For each intergenic region, we calculated a weighted score: the average binding site score of the two species * (1-nucleosome occupancy of *S. cerevisiae* promoter). Linear regression and Mann-Whitney tests were used to predict temperature effects by the weighted scores.

Binding site divergence for each binding site model was calculated by the difference between the highest scoring site for each allele. To test for associations between expression and the combined divergence of all binding sites we used the average of the absolute value of binding site divergence. For each motif model, linear regression was used to test association between binding site divergence and allele specific effects.

## Results

### Effects of temperature on allele-specific expression

To measure the effects of temperature on allele-specific expression (ASE) we generated RNA-seq data from an *S. cerevisiae* × *S. uvarum* hybrid during log phase growth at low (22°C), intermediate (33°C) and high (37°C) temperatures. Out of 5,055 orthologs, we found 2,950 (58%) that exhibited allele-specific expression, 1,669 (33%) that exhibited temperature-dependent expression and 136 (2.7%) that exhibited allele-by-temperature interactions (FDR < 0.05, Supplementary Data File). For the 1,669 temperature-responsive genes, expression levels were highly correlated between 33°C and 37°C (Pearson’s correlation coefficients = 0.97) and 37°C was more different from 22°C than 33°C (Pearson’s correlation coefficients = 0.89 for 22°C–37°C, 0.93 for 22°C–33°C). Despite the abundant temperature responses, allele differences were similar across temperatures with Pearson’s correlation coefficients of 0.90, 0.92 and 0.96 for 22–37°C, 22–33°C and 33–37°C, respectively (Figure 2A). In addition, the proportion of genes with the *S. cerevisiae* allele expressed at higher levels than the *S. uvarum* allele was 49.9, 50.7, 49.8% at 22, 33 and 37°C, respectively. Thus, there is no tendency toward higher *S. cerevisiae* allele expression at high temperature. *S. uvarum* alleles can be expressed at the same level as their *S. cerevisiae* ortholog at 37°C despite the fact that these promoters don’t experience high temperature in *S. uvarum* due to its temperature restriction.

Allele-specific temperature responses may reflect *cis*-regulatory changes involved in thermal differentiation. We therefore examined the 136 genes with a significant temperature-by-allele interaction. The gene set is not enriched for any GO terms (p > 0.05) and contains genes involved in a variety of biological processes. However, four genes are involved in trehalose metabolic process (*NTH2, TPS2, HSP104, PGM2*), and trehalose has been shown to influence thermotolerance (Eleutherio et al. 1993). Among the 136 genes, we found 27 where the *S. cerevisiae* allele responded to high temperature (37°C) more strongly than *S. uvarum*, and 40 genes where the *S. uvarum* allele responded more strongly. In the remaining 69 genes, alleles from the two species showed responses in opposite directions (Figure 2B).

**Fig. 2.**
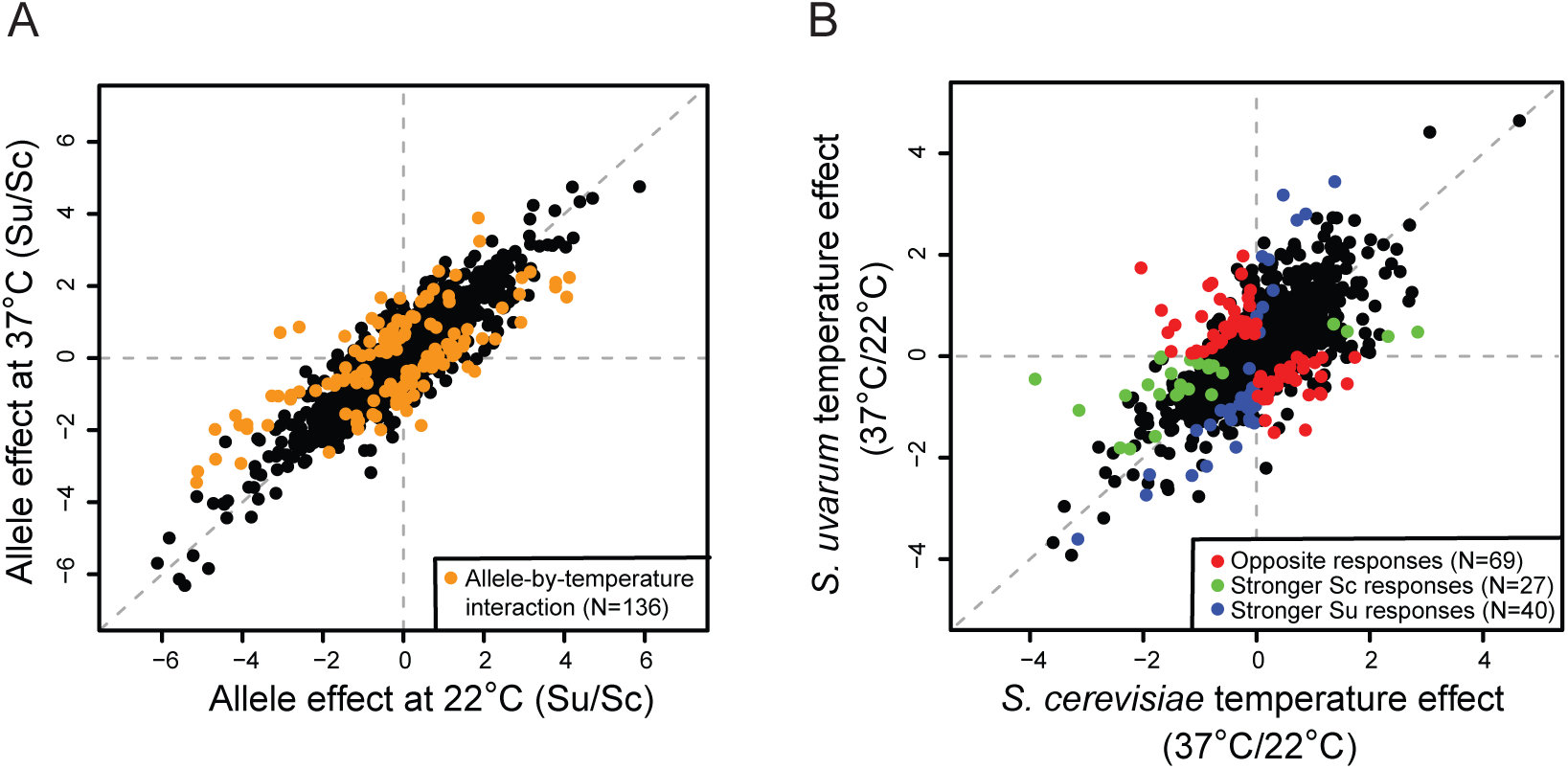
Temperature-dependent allele effects. The 136 genes with temperature-dependent allele effects are shown in color (legend) compared to all other genes (black, N = 4,919). A. Species’ allele effects (*S. uvarum/S. cerevisiae*) at low versus high temperature. B. Temperature effects (37°C/22°C) of *S. cerevisiae* (Sc) versus *S. uvarum* (Su). Temperature effects are classified into those with species’ alleles have an opposite temperature response (red), the *S. cerevisiae* allele responding to temperature more strongly than *S. uvarum* (green), and the *S. uvarum* allele responding to temperature more strongly than *S. cerevisiae* (blue).

### Effects of temperature on hybrid gene expression

To characterize temperature-dependent changes in gene expression we examined 211 genes that showed both a significant temperature effect (FDR < 0.05) and a 2-fold or more difference between the low (22°C) and high (37°C) temperatures. Unexpectedly, genes expressed at higher levels at the low temperature were enriched for genes involved in protein folding (*AHA1, MDJ1, BTN2, SSA2, HSP104, HSC82, SIS1, STI1, HSP82, CUR1*, p = 0.00829, Table S1). Typically, protein chaperones are induced in response to heat stress or misfolded proteins (Verghese et al. 2012).

To confirm the higher expression of genes involved in protein folding at 22°C and test whether this expression is specific to the hybrid or also found in one of the parents, we examined *HSP104* expression by quantitative PCR (Figure 3). Similar to our RNA-seq data, in the hybrid *HSP104* is expressed at higher levels at low temperatures (10°C and 23°C) compared to high temperatures (37°C) (5–fold change, p = 0.0006, t-test). Consistent with prior work (Gasch et al. 2000), in both parental species *HSP104* is expressed at the same level across temperatures and any transient induction that might have occurred upon a shift to 37°C is no longer present (linear regression, p = 0.11 for *S. cerevisiae* and 0.13 for *S. uvarum)*. However, in *S. uvarum HSP104* is expressed at higher levels than *S. cerevisiae* across all temperatures (t-test, p = 0.007, 0.013, 0.006 for 10°C, 23°C and 37°C, respectively). The atypical pattern of *HSP104* expression in the hybrid can be explained by a change in the dominant trans-acting environment. At low temperatures (10°C and 23°C) *S. uvarum* tends to dominate the trans-environment leading to high levels of *HSP104* expression whereas at 37°C *S. cerevisiae* completely dominates the *trans*-environment leading to low levels of *HSP104* expression.

**Fig. 3.**
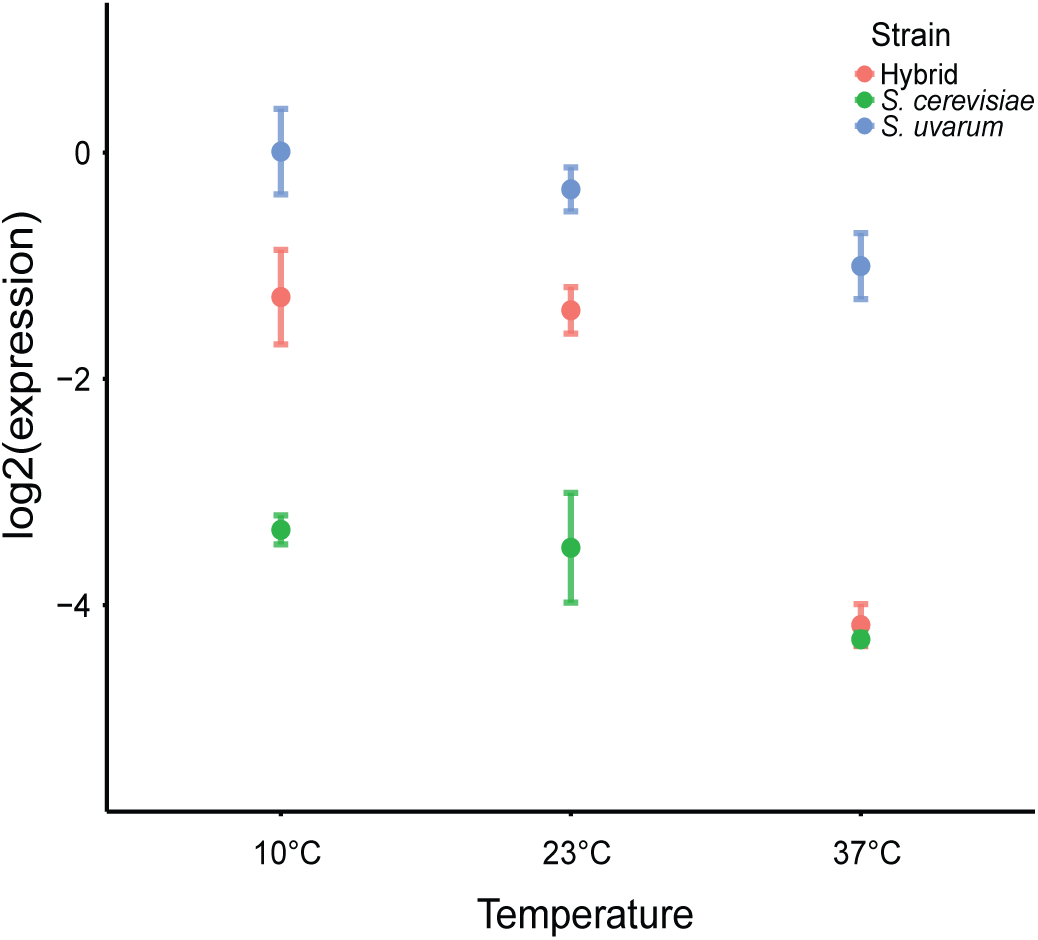
Temperature dependent *HSP104* expression in *S. cerevisiae, S. uvarum* and their hybrid. Expression is based on qPCR with points showing the mean and bars the standard errors. Hybrid expression is the sum of the two alleles.

### Test for temperature-specific directional evolution

Under a neutral model with no change in the selective constraints on gene expression, allele-specific differences in gene expression between species are expected to be symmetrically distributed. Parallel directional changes in gene expression among a group of functionally related or co-regulated genes can reflect selection (Bullard et al. 2010, Fraser 2011). Such groups have been reported in a hybrid of *S. cerevisiae* and *S. uvarum* by a sign test (Bullard et al. 2010), but the phenotypic consequences of these expression changes are not known. We tested whether patterns of directional selection are temperature-dependent, as might be expected if they are related to thermal differentiation. For example, consistent higher expression of the *S. cerevisiae* allele at the high but not low temperature would implicate directional selection in temperature-dependent expression divergence. We applied the sign test to ASE at each temperature separately and found 8, 9 and 13 groups of genes with directional ASE at 22°C, 33°C and 37°C, respectively (p < 0.01, FDR = 0.27, 0.24, 0.068 for 22°C, 33°C, 37C, respectively; Table 1, S2–S4). Seven groups are significant for all three temperatures, including the previously reported histidine biosynthesis and lysine biosynthesis groups (Bullard et al. 2010). Although we found a few groups specific to one or two temperatures using the p < 0.01 cutoff (e.g. Cluster_MET31, Cluster_adata– CalciumSpecific, etc.), all of these groups showed similar sum of scores across temperatures and p < 0.10 (Table 1). Therefore, none of the groups exhibiting directional divergence are temperature-specific.

**Table 1.** Groups of genes showing directional evolution at three temperatures.

| Group<sup>1</sup> | Number of genes in the group<sup>2</sup> | Sum of scores<sup>3</sup> |  |  | Annotation<sup>4</sup> |
| --- | --- | --- | --- | --- | --- |
|  |  | 22°C | 33°C | 37°C |  |
| Cluster_FHL1 | 93 (89) | 29** | 27** | 17* | Ribosomal proteins |
| Cluster_RPs | 136 (129) | 38** | 40** | 29** | Ribosomal proteins |
| Cluster_Histidine | 8 (6) | 6** | 5* | 4* | Histidine biosynthesis |
| Node 39 | 36 (24) | -12** | -10* | -8* | Organonitrogen catabolism |
| Cluster_MET31 | 17 (17) | 10* | 6 | 6* | Amino acid metabolism |
| Cluster_Lysine | 9 (9) | 6* | 7** | 5* | Lysine biosynthesis |
| Cluster_TFs | 18 (12) | -7* | -7* | -5* | Transcription factors |
| Node 81 | 7 (7) | 5* | 5* | 5** | Lysine biosynthesis |
| Cluster_adata-CalciumSpecific | 71 (44) | -10 | -15** | -15** | Membrane localization |
| Node 36 | 182 (144) | -20 | -17 | -19* | Unknown |
| Cluster_SWI6 | 29 (28) | -7 | -6 | -8* | Cell cycle regulation |
| Node 8 | 83 (64) | -10 | -7 | -11* | Oxidative stress |
| Cluster_PHO4 | 14 (12) | 5 | 5 | 5* | Unknown |
| Node 87 | 8 (5) | 4 | 4* | 3 | Microtubule polymerization |

<sup>1</sup> Groups are defined by Gasch et al. (2004). See Table S2-S4 for results of all groups.
<sup>2</sup> Number of genes with available data is shown in parentheses.
<sup>3</sup> Positive scores indicate *S. cerevisiae* alleles are expressed higher than *S. uvarum* alleles; negative scores indicate *S. cerevisiae* alleles are expressed lower than *S. uvarum* alleles. Significant groups are indicated for $p < 0.01$ (\*) and $p < 0.001$ (\*\*).
<sup>4</sup> The groups are annotated based on GO terms of genes in the group.

### Promoter changes associated with expression divergence

To identify promoter features that could explain allele-specific differences in expression we examined intergenic substitution rate, transcription factor binding site scores and their interaction with nucleosome occupancy. Among ASE genes, intergenic substitution rates were weakly correlated with gene expression divergence (Spearman’s rho = 0.064, p = 0.002). Given these differences we also calculated rates of binding site divergence using binding sites scores from 244 transcription factor binding site models (de Boer and Hughes 2012) and found a weak correlation between expression divergence and binding site divergence (Spearman’s rho = 0.05, p = 0.0119).

To identify binding sites that could explain allele-specific expression we first tested each binding site model for its ability to predict temperature responsive genes (22°C vs 37°C). We identified 17 motifs associated with genes induced at 22°C and 13 motifs associated with genes repressed at 22°C (Holm– Bonferroni corrected p < 0.05 for both linear regression and Mann-Whitney test, Figure S1). Many of the motifs (11/17) associated with up-regulated genes are similar to the stress response element (AGGGG), including the canonical stress response factors *MSN2* and *MSN4*. Other motifs known to be involved in the stress response include the heat shock factor *HSF1*, which is consistent with the observed up-regulation of heat shock genes at 22 °C (Table S1). Motifs enriched upstream of down-regulated genes are involved in glucose repression, e.g. *MIG1,MIG2, MIG3* and *ADR1*. *UME6* was also found, consistent with down-regulation of meiotic genes at 22°C revealed by GO analysis (Table S1). We also examined the correlations using a weighted score that accounts for both TF binding and nucleosome occupancy (see Methods), but the correlations were not greatly improved with the nucleosome weighted binding site scores.

Given the motifs associated with the temperature response, we tested each motif for an association between binding site divergence and ASE at 22°C. Within genes down-regulated at 22°C, divergence of 5 motifs was found to have a weak but significant association with expression divergence (*MIG1, MIG2, MIG3, TDA9* and *YGR067C*, linear regression, Holm-Bonferroni corrected p < 0.05, Figure S1). No motifs were correlated with ASE in genes up-regulated at 22°C, but one motif (*ARO80*) was correlated with ASE in genes that showed allele-by-temperature effects and up-regulation at 22°C (p = 0.0028, adjusted r-squared = 0.11). The weak correlations suggest that ASE is likely often caused by *cis*-regulatory mutations outside of known binding sites.

## Discussion

Environment-dependent gene expression is likely an important component of fitness. While *cis*-acting divergence on gene expression is abundant between species, the extent to which these *cis*-effects are environment-dependent is not often known. In this study, we show most *cis*-effects are independent of temperature in two thermally diverged yeast species. Further, we find that most *S. uvarum* alleles are expressed at levels similar to *S. cerevisiae* alleles at 37°C, even though *S. uvarum* does not grow at this temperature. Below, we discuss these results in relation to prior studies of variation in gene expression across environments and discuss the challenge of identifying changes in promoter sequences responsible for divergence in gene expression.

### Environment-dependent cis-effects

Changes in gene regulation may be an important aspect of how species adapt to different environments. Although there is extensive variation in gene expression-by-environment interactions (Hodgins-Davis and Townsend 2009), the extent to which these differences are caused by *cis*- or *trans*-acting factors is not as well characterized. We find that most *cis*-effects do not depend on temperature, only 136 of the 2,950 genes exhibiting ASE show temperature-dependent ASE. Thus, even though *S. uvarum* promoters have never been exposed to high temperatures, they can drive expression levels similar to those of *S. cerevisiae*. The consistent *cis*-effects across temperatures suggest that most *cis*-regulatory divergence is not associated with thermal divergence between the two species. Previous studies also found that *cis*-effects tend to be constant across environments and only a small subset of them are environment-dependent (Smith and Kruglyak 2008, Tirosh et al. 2009, He et al. 2016, Fear et al. 2016). Although we did not examine *trans*--effects genome-wide, the shift in the *trans*-effect of *HSP104* with temperature is consistent with prior work showing that *trans*-effects play a more pronounced role in environment-dependent differences in gene expression (Smith and Kruglyak 2008, Tirosh et al. 2009).

Although only a small number of genes showed a significant allele-by-temperature interaction, some may be relevant to thermal differentiation. Of particular interest are genes where the *S. cerevisiae* but not the *S. uvarum* allele responded to temperature. One noteworthy example of such is *TPS2*, which showed 2.5-compared to 1.5-fold induction of the *S. cerevisiae* compared to the *S. uvarum* allele, respectively. *TPS2* is involved in trehalose biosynthesis and essential to heat tolerance in *S. cerevisiae* (De Virgilio et al. 1993). The lower *cis*-regulatory activity of *TPS2* in *S. uvarum* might cause lower levels of trehalose and compromise its heat tolerance. In addition, three other genes (*NTH2, HSP104, PGM2*) in the trehalose pathway also showed allele-by-temperature effects, suggesting that the transcriptional regulation of this pathway might have diverged in the two species.

Among the 136 genes with temperature-dependent ASE, 67 genes showed a consistent direction but different magnitude of response for the two species’ alleles. The majority of them (53) were differentially induced at 22°C compared to 37°C and many are known to be induced by heat (*PIC2, SSE2, YKL151C, SIS1, IKS1, AHA1, EDC2, GSY2, HSP104, PUN1, TPS2*), oxidative stress (*ZWF1, YPR1, SOD1*) or other stresses (*CMK2*), consistent with the hybrid exhibiting a stress response at 22°C.

However, there is no bias for the *S. cerevisiae* or the *S. uvarum* allele being more induced (23 vs. 30 genes). In addition, in several heat-related genes (*AHA1, GSY2, HSP104*), the *S. cerevisiae* allele is more induced at 33°C than the *S. uvarum* allele, but at 22°C they are equally induced (with expression levels higher than or equal to those at 33°C). However, interpreting these changes is difficult given the *trans*-acting stress response is strongest at 22°C.

The 69 genes with alleles showing opposite responses to temperature are also worth discussing as some of them might be indicative of misregulation or thermal divergence. We examined their ASE pattern at 22°C and 37°C and classified them based on: ASE at both temperatures (24 genes), ASE at one temperature (42 genes), or ASE at neither of the two temperatures (3 genes). Among the 66 genes that showed ASE at one or more temperatures, only two genes (*IMP2’, POR2*) showed ASE at both temperatures but with opposite allele effects, where the *S. cerevisiae* alleles were higher than the *S. uvarum* alleles at 22°C but lower at 37°C. For the rest of the 64 genes, they either had ASE at both temperature but one allele consistently higher than the other allele, or showed ASE at one temperature but not the other. Thus, the 64 genes can be classified into two groups with 22°C-divergent or 37°C-divergent expression patterns. Two-thirds of them (43 genes) showed larger allele differences at 22°C than 37°C, i.e. 22°C-divergent. Among these genes, the *S. cerevisiae* alleles were expressed higher than the *S. uvarum* alleles at 22°C in 22 genes, vice versa for the remaining 21 genes. Interestingly, many genes in this group are related to mitochondrial function or oxidative stress (*GAD1, TIR3, QRI7, AIM41, YIG1, LAM4, YKL162C, THI73, ARG7, ICY1, YJL193W, YNL200C, YNL144C, YNL208W*). Mitochondrial function has been shown to be related to *S. cerevisiae’s* thermotolerance (Davidson and Schiestl 2001); thus the *cis*-acting divergence in mitochondria-related genes might be important to thermal divergence. In addition, the hybrid strain used in this study carries only *S. cerevisiae* mitochondrial DNA (mtDNA). Although it is also possible that the responses of mitochondria-localized genes are affected by *S. cerevisiae* mtDNA, this would imply species-specific feedback regulation on mRNA levels.

Besides the mitochondrial genes, membrane proteins (*YLR046C, YJR015W, THI73*), cell wall (*TIR3, CWP1*) and mating-related genes (*PRM4, AXL1, SIR1*) were also found in the 22°C-divergent group. The 21 genes in the 37°C-divergent group are involved in responses to glucose limitation (*GTT1, GSY1*), sporulation (*QDR3, NPP1*), cell signaling (*RHO5, TOS3, ROM1*), nutrient metabolism (*QDR3, YJR124C, NPP1, STR2, ATF2*) and mitochondrial functions (*TOS3*).

Taken together, one of the most notable features of the allele responses is that they more often diverge at 22°C than 37°C (43 vs. 21). Given that expression at 22°C resembles a stress response (Table S1), the greater divergence at 22°C may reflect divergent stress responses between the two species. Although the genes with allele-specific temperature responses have diverse biological functions, the stress- and mitochondrial-related genes are more often differentially induced at 22°C. However, it is also important to consider that these differences may only be present in a hybrid environment where we find a stronger stress response at low compared to high temperature.

### Unexpected heat shock response at low temperatures

The non-canonical expression of heat shock genes at 22°C is somewhat perplexing. Because we measured expression at constant temperatures we did not expect to see induction of heat shock genes, which normally occurs within 30 minutes of treatment and then dissipates (Gasch et al. 2000). Given the high expression level of *HSP104* in *S. uvarum* across all temperatures, one potential explanation for the heat shock response is a *trans*-signal produced by the *S. uvarum* genome. The absence of the heat shock response in the hybrid at high temperature may be a consequence of loss of the *S. uvarum trans*-signal, although this does not explain the high *HSP104* expression at high temperature in *S. uvarum*. Sample mix-up is unlikely as the *HSP104* experiment was done independently and is consistent with the original RNA-seq experiment.

The heat shock gene expression profile shows that the hybrid is under stress at 22°C but not 37°C. To better understand this counterintuitive phenomenon, we compared the hybrid expression profile to previously published *S. cerevisiae* (Gasch et al. 2000) and *S. uvarum* (Caudy et al. 2013) datasets. The hybrid temperature effect (37°C over 22°C) associates with 285 of 477 stress responses of either *S. cerevisiae* or *S. uvarum* (Spearman’s correlation test, Holm-Bonferroni corrected p < 0.05). However, 232 of the 285 correlations are negative, implying that 22°C is more stressful than 37°C in the hybrid. Interestingly, the strongest positive correlation is between the hybrid’s temperature response and *S. uvarum*’s 17°C to 30°C response at 60 min (Spearman’s rho = 0.23, Holm-Bonferroni corrected p = 5.39E-48). In contrast, the correlations with *S. uvarum*’s 25°C to 37°C or 25°C to 42°C response are negative. Similar to the hybrid, heat shock genes are expressed higher at 17°C than 30°C in *S. uvarum*, but the pattern is not seen in the other two temperature shifts (Caudy et al. 2013). These differential correlations indicate *S. uvarum*’s heat shock response may be sensitive to specific temperatures used in the shifts. However, it is also important to note that heat shock proteins are not specific to temperature changes but are part of the general environmental stress response which can be induced by any number of environmental changes (Gasch et al. 2000). Taken together, the stress response induced in the hybrid at 22°C may reflect a contribution from the non-canonical temperature response in *S. uvarum*.

### Signatures of selection on cis-acting divergence in gene expression

The sign test of allele imbalance across functionally related genes has been used in a variety of configurations to detect polygenic *cis*-regulatory adaptation (Bullard et al. 2010, Fraser et al. 2010, Fraser 2011, Naranjo et al. 2015, He et al. 2016). However, previous applications of the test were to expression levels under standard growth conditions. Because gene expression is environment-dependent, some signals of selection may only be uncovered by examining expression in environments to which an organism adapted. However, our results indicate that directional ASE as found by the sign test is not temperature-dependent, consistent with our observation that most *cis*-effects are not temperature dependent. Our results do not rule out the possibility of *trans*-acting expression differences important to thermal differentiation, nor do they address *cis*-acting changes that occur immediate after a temperature shift and which are typically much stronger and more widespread than those that persist for hours after the initial shift (Gasch et al. 2000).

In addition to the histidine and lysine biosynthesis groups reported by Bullard et al. (2010), we found several other groups of genes showing a signature of directional evolution. Among these, the ribosomal genes show a strong bias toward higher *S. ceverisiae* allele expression (Table 1), which could indicate a difference in translational capacity of the two species. Two other groups, Node 39 (organonitrogen catabolism) and Cluster_MET31 (amino acid metabolism) provide new evidence for divergence in nutrient metabolism between the two species.

Most groups identified by the sign test contain a substantial number of temperature-responsive genes, with the lysine biosynthesis pathway showing the highest fraction (8/9). The pathway consists of nine genes (*LYS1, LYS2, LYS4, LYS5, LYS9, LYS12, LYS14, LYS20, LYS21*), eight of which are induced at 22°C, with *LYS4* and *LYS20* showing allele-by-temperature effects. The *S. cerevisiae* allele of *LYS20* is induced at 22°C more than the *S. uvarum* allele (3.2 vs. 1.0 fold). Although not a significant temperature-by-allele interaction, a similar pattern is present for *LYS1* (4.1 vs. 2.9 fold) and *LYS2* (2.3 vs. 2.1 fold).

The weak responses of *S. uvarum* alleles might reflect deficiency in activating the lysine biosynthesis pathway at a given temperature or under stress, which is critical for amino acid homeostasis. Also, the lysine biosynthesis pathway is known to be induced by mitochondrial retrograde signaling in response to compromised mitochondrial respiratory function (Liu and Butow, 2006) and could potentially be affected by the *S. cerevisiae* mtDNA.

### Binding sites are only weakly related to expression divergence

Consistent with previous reports (Tirosh et al. 2008;Tirosh and Barkai 2008;Chen et al. 2010;Zeevi et al. 2014), we only found weak correlations between binding site changes and allele-specific expression. Previous work has shown that binding sites in nucleosome depleted regions are more likely to cause changes in gene expression (Swamy et al. 2011). Yet, incorporation of predicted nucleosome occupancy did not improve our ability to predict gene expression. This finding is consistent with another study that found no relationship between divergence in nucleosome occupancy and gene expression in yeast (Tirosh et al. 2010). One explanation for the weak correlations is that ASE may often be caused by *cis*-regulatory mutations outside major binding sites, e.g.Levo et al. (2015). Genes in the lysine biosynthesis pathway provide a good example of conserved binding sites: seven genes in the pathway showed higher *S. cerevisiae* expression, yet binding sites for *LYS14*, the major transcription factor that regulates these genes (Becker et al. 1998), are conserved in all of them. Furthermore, the lysine genes are also not enriched for divergence in other motifs present upstream of these genes (e.g. *MOT2, XBP2, RTG1, RTG3*, p > 0.05, Mann-Whitney test).

Despite binding site divergence being only weakly related to ASE, we found a few significant associations with specific binding sites. One of these, *ARO80* sites, correlated with temperature-dependent expression differences largely due to two genes *ARO9* and *ARO10* (Figure S1, S2). In both cases, the *S. uvarum* promoters have lower binding scores and lower expression of the *S. uvarum* allele (Figure S2). Interestingly, the number of monomers in the *ARO80* binding sites also differs between *S. cerevisiae* and *S. uvarum*. In both genes, *S. cerevisiae* sites are tetrameric and *S. uvarum* sites are trimeric (Figure S2). The example of *ARO80* suggests expression divergence might associate with changes in the number of binding sites, which our binding site analysis didn’t consider.

## Acknowledgements

We thank Chris Hittinger for sharing the *S. uvarum* strain, Ching-Hua Shih for help with the RNA-seq analysis pipeline and other members of Fay lab for constructive comments. This work was supported by an NIH grant (GM080669) to JCF.

